# DNA Topology in Chromatin is Defined by Nucleosome Spacing

**DOI:** 10.1101/104083

**Authors:** Tatiana Nikitina, Davood Norouzi, Sergei A. Grigoryev, Victor B. Zhurkin

**Affiliations:** Laboratory of Cell Biology, CCR, National Cancer Institute NIH, Bethesda, MD, USA; Department of Biochemistry and Molecular Biology, Pennsylvania State University College of Medicine Hershey, PA, USA

## Abstract

In eukaryotic chromatin, DNA makes about 1.7 left superhelical turns around an octamer of core histones implying that formation of nucleosomes would alter the overall topology of DNA by a comparable difference of the DNA linking number (Δ*Lk*) per nucleosome. However, earlier experiments have documented a significantly (about 50%) lower absolute value |Δ*Lk*| than expected from the nucleosome geometry. Recently, using computer modeling, we have predicted two families of energetically stable conformations of the arrays with precisely positioned nucleosomes, one with an integer number of DNA turns in the linker DNA {L = 10*n*} and the other with extra five base pairs in the linker {L = 10*n* + 5}, to be topologically different. Here, using arrays of precisely positioned clone 601 nucleosomes, topological electrophoretic assays, and electron microscopy we experimentally tested these predictions. First, for small 12-mer nucleosome circular arrays we observed that dLk per nucleosome changes from −1.4 to −0.9 for the linkers {L = 10*n*} and {L = 10*n* + 5}, respectively. Second, for larger hybrid arrays containing a mixture of positioned and non-positioned nucleosomes we found that changing the DNA linker length within the positioned arrays was sufficient to significantly alter the overall DNA topology fully consistent with our prediction. The observed topological polymorphism of the circular nucleosome arrays provides a simple explanation for the DNA topology in native chromatin with variable DNA linker length. Furthermore, our results may reflect a more general tendency of chromosomal domains containing active or repressed genes to acquire different nucleosome spacing to retain topologically distinct higher-order structures.

## INTRODUCTION

In eukaryotic chromatin, the DNA double helix is repeatedly supercoiled in the nucleoprotein particles called nucleosomes. The core of a typical nucleosome contains 145-147 bp of DNA making 1.7 left superhelical turns around a histone octamer (1-3). The nucleosomes play an essential role in regulating all DNA-dependent processes such as transcription, replication, recombination, and repair through local and dynamic unfolding of chromatin (4-6).

In the cell nucleus, the nucleosome cores are connected by relatively extended DNA linkers forming beads-on-a string nucleosome arrays. The nucleosome arrays fold into higher order structures that mediate DNA packing and availability for the DNA-recognizing machinery (7, 8). Importantly, transcriptionally-active chromosomal domains accumulate unconstrained negative supercoiling (9, 10) indicating that active chromatin may have different linear distribution of torsional stress than condensed heterochromatin. These findings necessitate experiments aimed to relate the intrinsic chromatin structural variations predicted *in vitro* and *in silico*, to DNA topologies and associated functional interactions *in vivo*.

Monitoring of DNA supercoiling in circular nucleosome arrays (minichromosomes) by topological gel assays has been long used as a way to analyze topological properties of DNA in chromatin. These assays are based on the helical periodicity of DNA making one turn per ~10.5 bp at physiological conditions (11, 12), which corresponds to helical Twist ≈ 34.5°. When the ends of the DNA chains are constrained in covalently closed circular DNA (ccDNA), the changes in DNA Twist (*Tw*) are compensated by the changes in Writhe (*Wr*) according to the well-known equation Δ*Lk* = Δ*Tw* + *Wr*, where the linking number (*Lk*) is defined as the number of times one strand of the duplex turns around the other (13-16). Note that the three parameters describing the DNA topology (Δ*Lk*, Δ*Tw* and *Wr*) are measured relative to the relaxed state of DNA. Therefore, Δ*Wr* = *Wr*, because the writhe of unconstrained DNA is zero.

For a chain of nucleosomes, the X-ray crystal structure of the nucleosome core (1-3) predicts generation of DNA supercoiling with *Wr* = Δ*Lk* = −1.7 per nucleosome if the DNA twist is not changed (Δ*Tw* = 0). However, multiple measurements of Δ*Lk* in ccDNA from either native minichromosomes of SV40 virus (17, 18) or nucleosome arrays reconstituted from histones and circular DNA (19) persistently resulted in experimental values of Δ*Lk* = −1.0. This discrepancy is known as the linking number paradox (16, 20-22). (Note that when acetylated histones were used for reconstitution, the linking number changed insignificantly, to Δ*Lk* = −0.8 (23).)

How could the nucleosome structure and topological properties of the nucleosome chain be reconciled? One explanation is that the DNA Twist in the nucleosome cores could differ from that of naked DNA in solution (20-22, 24, 25). The other possibility is that a special path of linker DNA could alter *Wr* (15, 26). Specific linker DNA geometries consistent with the observed DNA topology have been proposed (26-30). However, the actual linker DNA path within topologically constrained chromatin domains remained obscure due to the absence of adequate circular minichromosome models with precise nucleosome positions.

During the past dozen years there has been a significant progress in the structural studies of chromatin, largely due to a wide usage of strongly positioned ‘601’ nucleosome sequence (31). In particular, the seminal studies of Richmond and colleagues showed the two-start zigzag fibers in solution (32) and in crystal form (33). Their results, together with the high-resolution Cryo-EM data obtained by Song et al. (34) strongly support the two-start zigzag organization of chromatin fibers for relatively short nucleosome repeat lengths, NRL = 167, 177 and 187 bp. In addition, the electron microscopy (EM) images presented by Rhodes and co-workers confirmed zigzag organization for NRL = 167 bp, while for NRL = 197 bp and longer, more tightly packed fibers were observed, especially in the presence of linker histones (35).

All these structures were obtained for arrays of strongly positioned ‘601’ nucleosomes (31), with NRL varying from 167 to 237 bp in increments of 10 bp. Provided that the nucleosome core is 147 bp (2), the linker length L varies from 20 to 90 bp – that is, L belongs to the {10*n*} series. On the other hand, it is known that *in vivo*, the linker sizes are close to {10*n*+5} values (36-39). Thus, the structural data mentioned above correspond to linker lengths with occurrences *in vivo* that are relatively small.

Recently, we have analyzed positioned nucleosomal arrays with linker lengths belonging to the {10*n*} and {10*n*+5} series both experimentally and theoretically. Using analytical ultracentrifugation and EM imaging, we demonstrated that fibers with L = 25 bp have less propensity to fold in a compact state compared to L = 20 or 30 bp (40). Further, using computer modeling, we predicted the two families of energetically stable conformations of the two-start zigzag fibers, one with L = 10*n* and the other with L=10*n* + 5 to be topologically different, the former one imposing ~50% higher absolute |Δ*Lk*| values in the circular DNA (41, 42), see Fig. 1. Here, we experimentally addressed these predictions for circular nucleosome arrays containing regularly spaced repeats of the ‘601’ nucleosome positioning sequence (31) using 1D and 2D topological electrophoretic assays and electron microscopy. Our results are in excellent agreement with the predictions, thus demonstrating that topological polymorphism of the nucleosome arrays is largely defined by the inter-nucleosome linker lengths. These findings may reflect a universal tendency of chromosomal domains containing active or repressed genes to acquire different nucleosome spacing to retain topologically distinct higher-order structures.

**Fig. 1.**
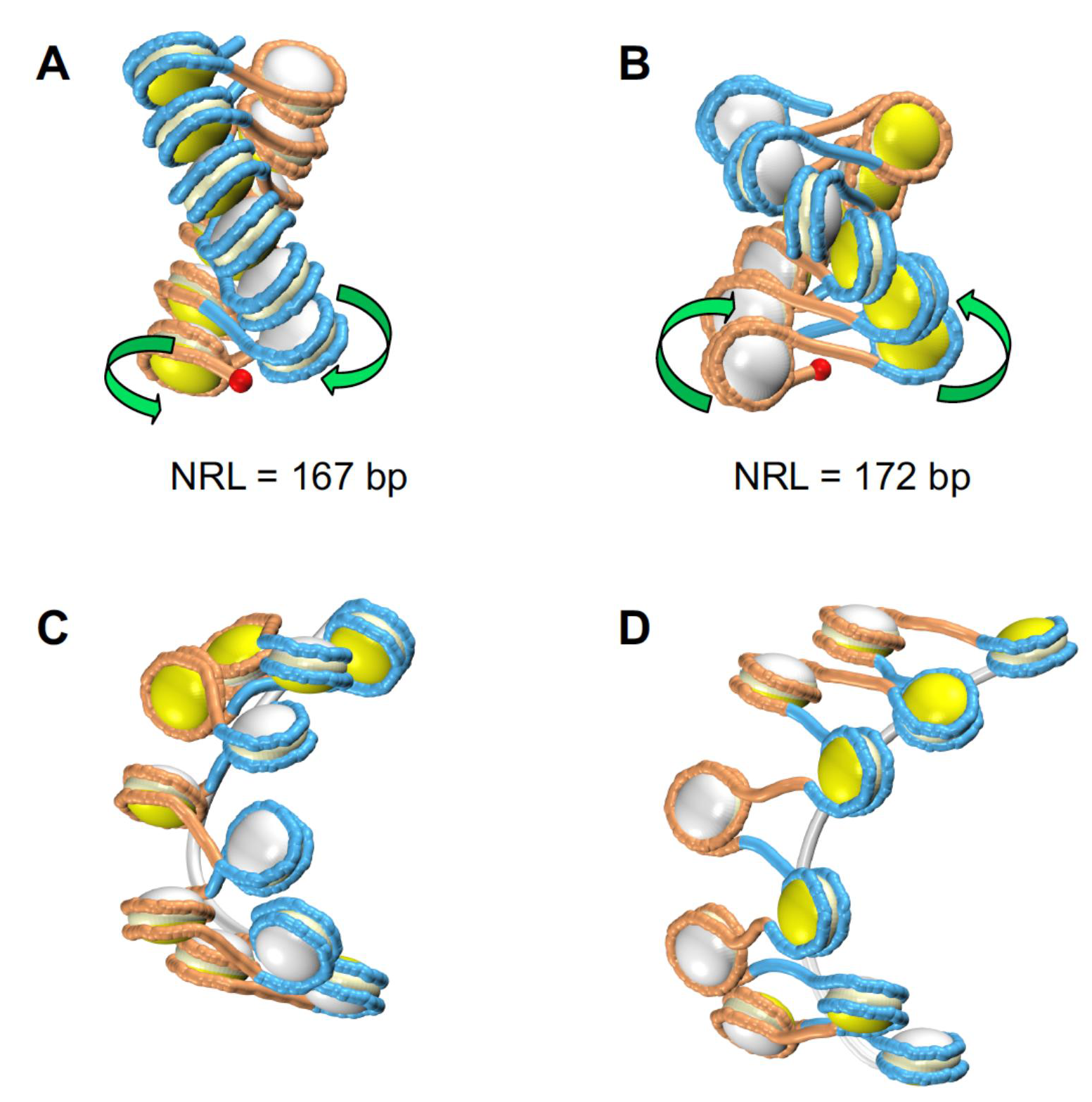
Models of two-start chromatin fibers with NRL=167 and 172 bp. (A, B) The energy-optimized regular fibers (41, 42) containing 12 nucleosomes, with NRL = 167 bp (A) and NRL = 172 bp (B). The DNA linking number per nucleosome, Δ*Lk* = −1.37 and −0.93, respectively. The DNA is presented in alternating blue and orange colors, to emphasize the two stacks of nucleosomes; the DNA ‘entry’ points are shown as red balls. The histone cores are shown in two colors—the ‘entry’ sides are in yellow, and the ‘exit’ sides in white. In this way, it is easier to distinguish the fiber configurations. In addition, the green arrows indicate different DNA folding pathways in the two topoisomers. (C, D) Representative configurations of the 167x11 and 172x11 circular nucleosomal arrays obtained in the course of energy minimization. (See main text and Fig. 4 for details.) Note that the circular topoisomers (C, D) are significantly distorted and extended compared to the regular conformers (A, B). The nucleosome-free DNA fragments are shown as white tubes.

## RESULTS

### Plasmid-based DNA circles

We prepared circular DNA templates using two different methods. First, we constructed plasmid-based DNA circles containing 12 repeats the ‘601’ sequence with NRL=167 or 172 bp inserted into pUC19 vector (see Methods for details). In these 4.7 kb-long circles, about half of DNA belongs to pUC19 vector, which is not a nucleosome positioning sequence. To ensure that presence of vector DNA does not affect the resulting measurements of Δ*Lk* difference, we also prepared DNA mini-circles consisting purely of 601 repeats (see below).

The plasmid-based circular constructs (denoted p-167x12 and p-172x12) were prepared by standard methods of cells transformation, DNA extraction and purification (see Methods for details). Plasmid DNA extracted from cells has superhelical density σ ≈ −0.06 (43), which is expected to facilitate formation of nucleosomes. Therefore, we used these plasmid-based circles for reconstitution of nucleosomes by a standard salt dialysis method. Examination of the resulting nucleosomal arrays (NAs) by electrophoretic mobility retardation assay on native agarose gel (Fig. S1) shows practically complete incorporation of DNA into the nucleosome bands at the 100% loading of the core histones. We also digested the NAs with restriction enzymes to mononucleosomes and verified the equal core histone loading by the nucleosome band shift assay (Fig. S2).

Next, DNA in the reconstituted nucleosomal arrays was relaxed with Topoisomerase I, deproteinized, and run on agarose gels containing different concentrations of intercalator chloroquine (CQ). Comparison of the electrophoretic mobility of scDNA obtained with variable histone loading allows monitoring increase in the superhelical density of DNA that accompanies formation of nucleosomes (Fig. 2). Our approach is illustrated in Supplementary Fig. S3.

**Fig. 2.**
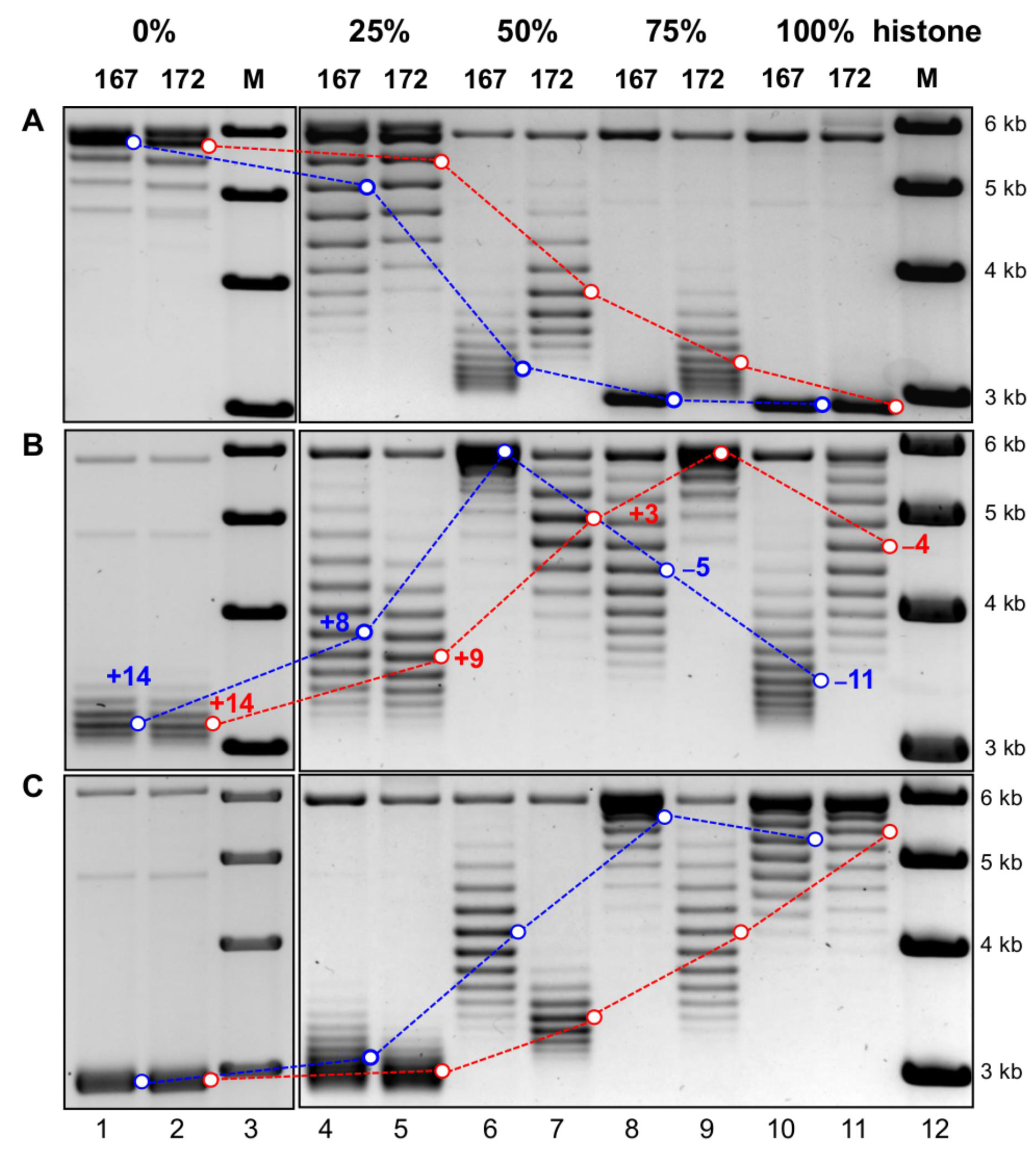
Topological difference between nucleosomal arrays with 167 and 172 bp NRL reconstituted on plasmid DNA. Plasmid-based circular DNA templates p-167x12 (lanes 1, 4, 6, 8, 10) and p-172x12 (lanes 2, 5, 7, 9, 11) were reconstituted with 0%, 25%, 50%, 75%, and 100% core histones (shown on the top), treated with Topoisomerase I, the circular DNA was isolated and separated on agarose gels run in the absence (A) and presence of 1.5 (B) and 4.0 (C) μg/ml CQ in the gel and TAE buffer. Lanes 3 and 12: molecular weight markers. The strongest bands are indicated by blue circles for the 167 bp NRL, and by red circles for 172 bp NRL. The numbers of superhelical turns in the topoisomers corresponding to these bands are given in the same colors (panel B). In particular, for the bands denoted +8 and +9 (lanes 4 and 5), the shifts from the top of gel (nicked circles, form II) are counted directly. For the bands denoted +14 (lanes 1 and 2), the shift from the top is evaluated from the relative shift of 6 supercoils between these bands and the band +8 (lane 4). Difference in DNA linking number between the 167 and 172 bp NRL constructs, or ∆(∆Lk), equals 1 at 25%, 5 at 50%, 6 at 75% and 7 at 100% loading of core histones. Note that the presence of intercalator CQ introduces additional positive supercoils in DNA and changes its electrophoretic mobility. For example, the DNA circles relaxed in the absence of histones (lanes 1 and 2), become strongly supercoiled and run at the bottom of gel. By contrast, the strong negative superhelical density of DNA obtained at 100% loading of histones (lanes 10 and 11) is partially compensated by CQ and thus, its mobility is decreased. (See Suppl. Fig. S3 for details.)

As follows from Fig. 2 A (lanes 1 and 2), both DNA circles, p-167x12 and p-172x12 (without core histones) are relaxed and run at the top of the gel. Gradual increase in the core histone loading increases the superhelical density and accordingly changes the electrophoretic mobility, entirely consistent with the scheme presented in Fig. S3. Specifically, the DNA samples obtained with 25% loading of histones have a ‘modest’ superhelical density (Figs. 2 A, lanes 4 and 5), which is increased at 50-75% loading, as reflected in a relatively high gel mobility (lanes 6-9). Finally, the 100% loading of core histones further increases superhelical density of DNA, so that the tightly coiled DNA migrates to the very bottom of gel and forms zones of poorly resolved bands (Fig. 2 A, lanes 10 and 11).

To resolve individual, fast migrating bands we repeated the electrophoresis in the presence of various concentrations of CQ, which is known to induce additional positive supercoils in circular DNA (44). The DNA circles obtained with a small loading of histones (0 and 25%), become positively supercoiled and run at the bottom of the gel (Fig. 2 B, lanes 1, 2; Fig. 2 C, lanes 1, 2, 4, 5). The DNA purified from NAs with 50-75% loading has intermediate superhelical density; its gel mobility is reversed in the presence of CQ (Fig. 2 B, lanes 6, 7; Fig. 2 C, lanes 6-9), in agreement with published data (45). And most importantly, for DNA with the highest superhelical density (with 100% histone loading) in the presence of 1.5 μg/ml CQ we observed a strong decrease in mobility so that individual bands can be resolved on the gel (Fig. 2 B, lanes 10 and 11).

Now, using gels presented in Fig. 2 B ([CQ] = 1.5 μg/ml), we can accurately determine the change in the DNA linking number induced by formation of nucleosomes. Based on the analysis presented above, we conclude that for the low histone loading (0 and 25%), the p-167x12 and p-172x12 constructs have similar number of positive superhelical turns. The dominant topoisomers are located either at the same position, band +14 (lanes 1, 2), or they are shifted by one band (lanes 4, 5; bands +8 and +9). For the 100% loading, both DNA circles, p-167x12 and p-172x12, have negative superhelical turns and now show a striking difference between the two samples. Evaluation of the linking number difference between the p-167x12 and p-172x12 nucleosome arrays is straightforward and gives a change of *Lk* = 7 negative supercoils in 167-bp *vs* 172-bp arrays. In the cases of 50 and 75% loadings (lanes 6 and 9), the strongest bands are located too close to the top of gels, and it is impossible to decide whether they correspond to the negative or positive supercoils from the 1D gels.

To resolve all topoisomers on a single gel, we run 2D electrophoreses (Fig. 3). The CQ concentrations were selected so that the strongest bands for the p-172x12 construct would be located at the top of gel, and for the p-167x12 construct they would be on the left side of gel. As a result, we see that the dominant topoisomers in the two constructs are shifted by 5-6 sc for 50% histones loading (Fig. 3 A) and by 6-7 sc for 75% loading (Fig. 3 B). Thus, we conclude that the topological difference between the p-167x12 and p-172x12, ΔΔ*Lk*, monotonically increases reaching about 7 supercoils, when the loading of core histones increases to 100%.

**Fig. 3.**
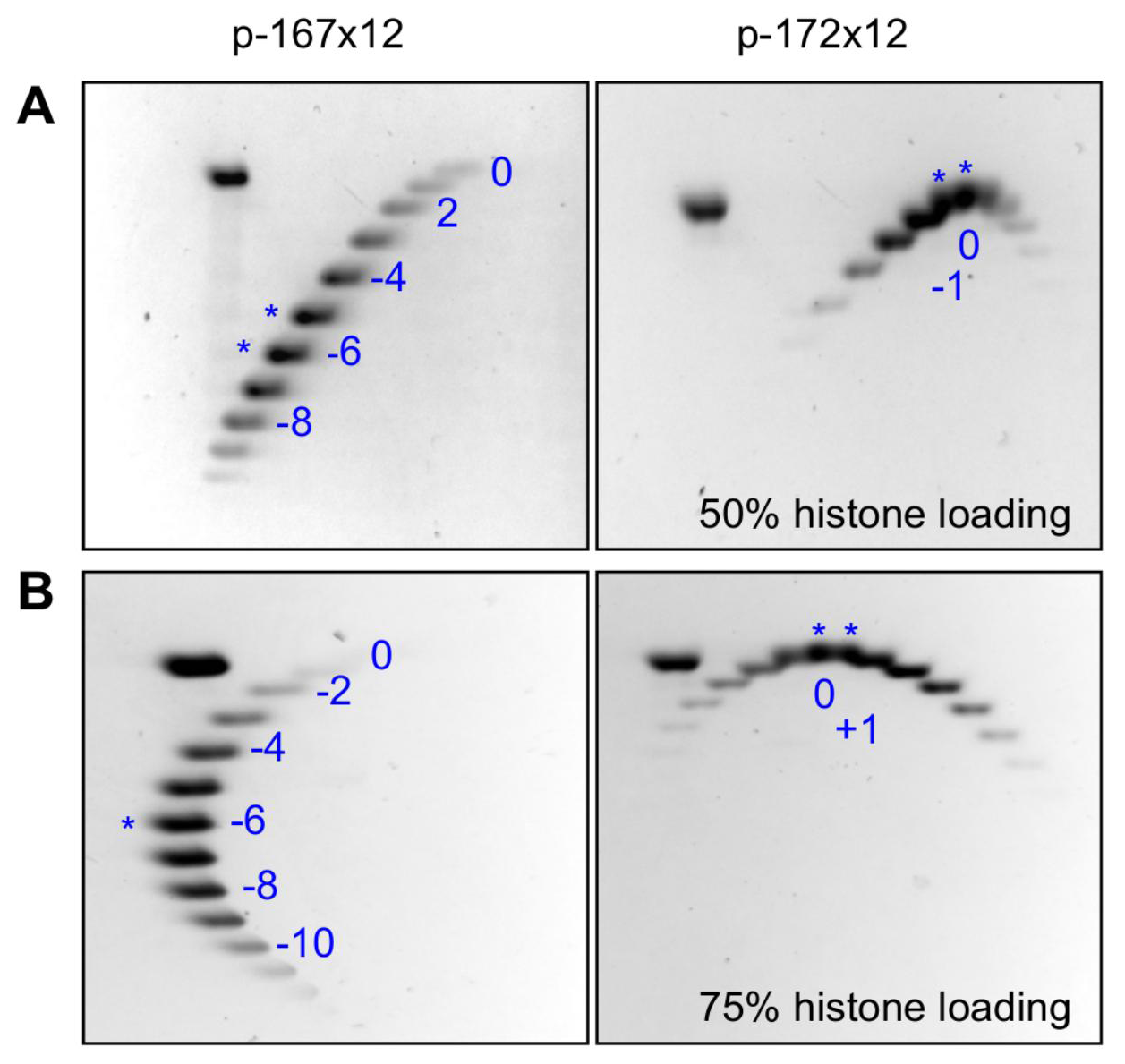
Two-dimensional gel electrophoresis of DNA extracted from nucleosomal arrays with 167 and 172 bp NRL. Plasmid-based circular DNA templates p-167x12 (left gels) and p-172x12 (right gels) were reconstituted with 50% (A) and 75% (B) core histones, treated with Topoisomerase I, the circular DNA was isolated, and separated in the first direction (downward) in the presence of 0.5 μg/ml CQ, and in the second direction (to the right) in the presence of 3.0 μg/ml CQ (A), or with 1.5 μg/ml CQ in the first and 4.0 μg/ml CQ in the second direction (B). Note that the dominant topoisomers with 167 bp NRL (left gels, marked by asterisks) have additionally 5-6 negative supercoils for 50% histone loading (A) and 6-7 negative supercoils for 75% loading (B) as compared with the 172 bp NRL (right gels).

Provided that the nucleosome arrays have the same number of ‘601’ nucleosomes and the vector-associated nucleosomes are not supposed to change between the two samples, our data are consistent with the model-predicted (Fig. 1) difference in ΔΔ*Lk* = 0.5 (per ‘601’ nucleosome) between the p-167x12 and p-172x12 constructs.

## DNA mini-circles

To make sure that the *Lk* difference between the two types of chromatin fibers observed above is not affected by the presence of vector DNA, we analyzed DNA mini-circles containing only the ‘601’ repeats. This approach is similar to that previously used in construction of the mini-circles based on sea urchin 5S DNA nucleosome positioning sequences by Simpson et al. (19) and Norton et al. (23). The 2 kb-long supercoiled DNA circles were prepared by ligation and relaxation by TopoI in the presence of various concentrations of EtBr (see Methods), which allowed producing scDNA with defined superhelical density (46). Gradually increasing concentration of EtBr up to 2.0 μg/ml and comparing electrophoretic mobility of scDNA in the presence of variable amounts of CQ, we were able to modulate the number of supercoils in each sample (Fig. S4). In particular, the scDNA obtained in the presence of 2.0 μg/ml EtBr has 12 negative supercoils (Fig. S4 D), which corresponds to superhelical density σ= −0.06 in the 2 kb-long mini-circle. (Note that this superhelical density is comparable to that in native *E. coli* plasmids (43).) The 167x12 and 172x12 mini-circles produced the same superhelical density (172x12 bands are shown by asterisks in Fig. S4). We used 167x12 scDNA as a reference for measuring the number of DNA supercoils in scDNA extracted from the nucleosomal arrays.

Reconstitution of nucleosomes on the scDNA formed in the presence of 2.0 μg/ml EtBr gave a result very similar to that in the case of the plasmid-based circles – namely, DNA in 167x12 nucleosomal arrays is more negatively supercoiled than in 172x12 arrays (Figs. 4 A and B). However, the topological difference (Δ*Lk*) between the two arrays is only 3 supercoils. Also, the number of DNA supercoils in these arrays (11 and 8 sc respectively) is less than expected for complete nucleosome saturation (Figs. 4 A and B). This result is consistent with the number of nucleosomes directly counted on the EM micrographs (Fig. 4 C, D and supporting pdf file). Instead of 12 nucleosomes expected for the 100% loading of core histones, only about 9 nucleosomes on average are formed on both constructs, 167x12 and 172x12 (Figs. 4 C, D). Most likely, the limited number of nucleosomes formed on the 2 kb-long circles is due to the decreased conformational flexibility of DNA during nucleosome reconstitution in the mini-circles (compared to the 4.7 kb-long plasmid-based circles).

**Fig. 4.**
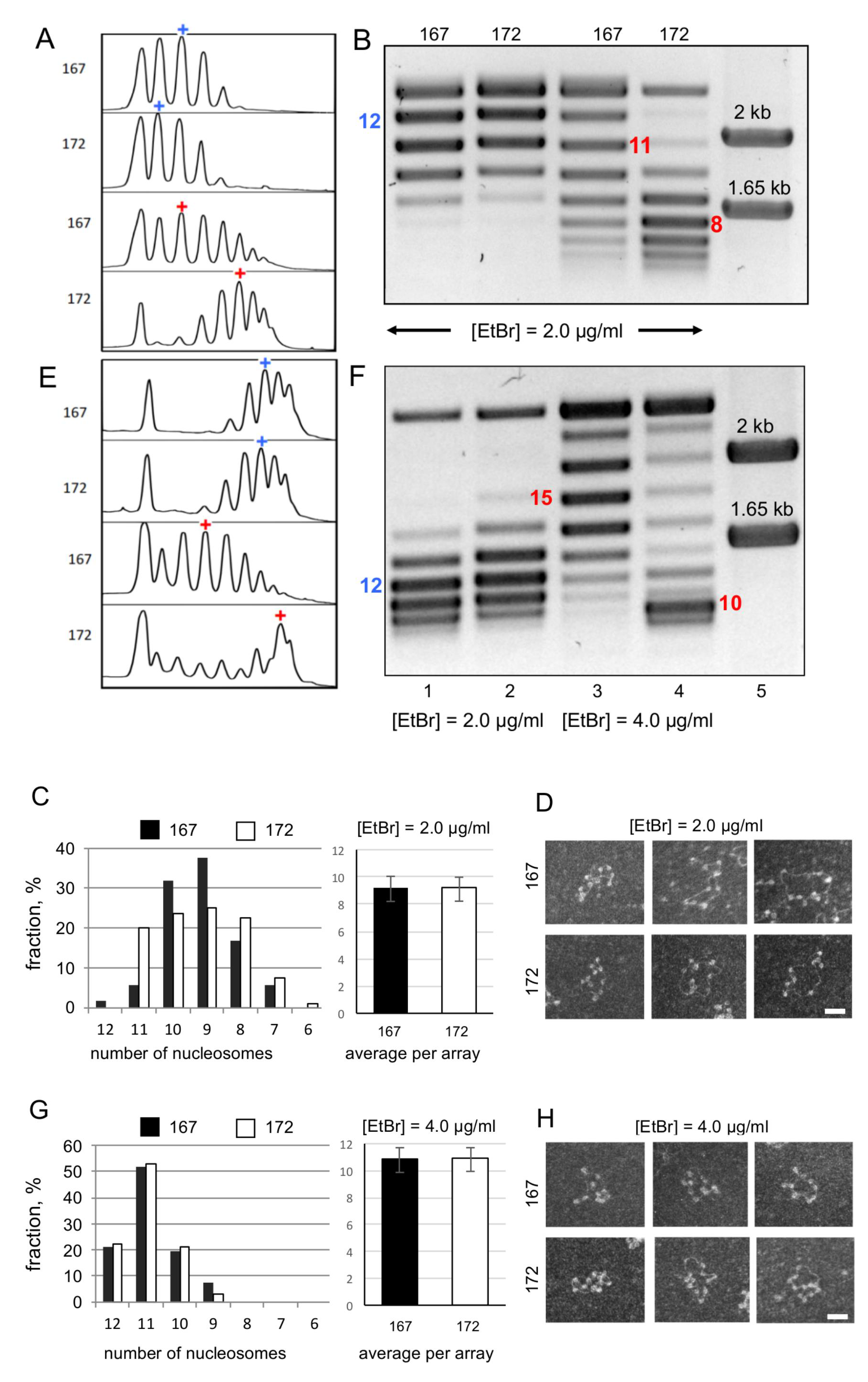
Topological difference between 167x12 and 172x12 nucleosomal arrays reconstituted on mini-circular DNA. Nucleosomes were reconstituted on the scDNA templates prepared in the presence of 2 μg/ml EtBr (A-D) and 4 μg/ml EtBr (E-H). A and E: Scans of gels in B and F. The strongest bands are marked with “+”. B and F: Electrophoretic mobility of the DNA topoisomers extracted from nucleosomal arrays was compared with the mobility of scDNAs templates obtained at [EtBr]= μg/ml (lanes 1 and 2). Gels were run in the presence of 8 (B) and 32 (F) μg/ml CQ. Lanes 5 show molecular markers. Note that the scDNAs with 167 bp (lanes 1) and 172 bp NRL (lanes 2) have similar distributions of topoisomers, with the strongest bands corresponding to *Lk* = −2 (Fig. S4 D). Nucleosomal arrays reconstituted on these scDNAs (obtained at [EtBr]=2 μg/ml) have *Lk* = −11 and −8 (lanes 3, 4, panel B). Arrays reconstituted on scDNA obtained at [EtBr]=4 μg/ml have *Lk* = −15 and −10 (lanes 3, 4, panel F). C and G: graphs showing the average number of nucleosomes on 167x12 and 172x12 arrays and distributions of the number of nucleosomes per one mini-circle calculated from EM images (Supporting pdf file). D and H: Representative transmission EM images of relaxed mini-circular 167x12 and 172x12 arrays reconstituted on scDNA obtained at [EtBr]=2 μg/ml (D) and [EtBr]=4 μg/ml (H). Scale bar 50 nm.

To facilitate formation of nucleosomes, we increased the number of negative supercoils in mini-circular DNA, using scDNA prepared in the presence of 4.0 μg/ml EtBr (Fig. S5). As a result, the number of nucleosomes detectable on the EM micrographs increased to ~11 on average, the distribution of the corresponding numbers narrowed, and the mini-circles containing the full amount of 12 nucleosomes became rather abundant (Fig. 4 G, H). Accordingly, the number of supercoils increased up to 15 and 10 for NRL=167 and 172 bp, respectively (Figs. 4 E and F). For illustration, the representative conformations of 167x11 and 172x11 NAs obtained in the course of energy minimization are shown in Figs. 1 C, D.

Note that the average numbers of nucleosomes in the 167 bp NRL arrays (10.86) and in the 172 bp NRL arrays (10.94) are virtually equal (Fig. 4 G), therefore we can exclude the possibility that the stronger superhelical density observed for NRL = 167 bp is the result of the higher histone loading in this case. Thus, the topological difference between the uniform 10*n* and 10*n*+5 nucleosomal arrays increased to 5 supercoils, or 45% of the total number of nucleosomes, in excellent agreement with our model (Fig. 1) predicting the dependence of the nucleosome array folding and topology on the linker DNA length.

## DISCUSSION

Our results reconcile the nucleosome core crystal structure with topology of the nucleosome array. For the first time, we show that the level of DNA supercoiling in the nucleosomal arrays varies by as much as ~50% depending on the DNA linker length. The DNA linking number per nucleosome, Δ*Lk*, changes from −1.4 for L = 20 bp to −0.9 for L = 25 bp (in DNA mini-circles consisting purely of ‘601’ repeats). Because the same sequence ‘601’ was used in both cases, one can be certain that the observed change in DNA supercoiling is due to changes in the global DNA pathway (Writhe) rather than to alteration of DNA twisting in nucleosomes. Note that the nucleosome positioning signals embedded in this sequence are so strong that they secure the same positioning of the ‘601’ nucleosomes both in array and in a single nucleosome with a single nucleotide precision (47, 48). Furthermore, changing the DNA linker length in the ‘601’ repeats is sufficient to significantly alter the overall DNA topology in hybrid plasmids containing a mixture of positioned and non-positioned nucleosomes and thus mimicking native nucleosomal arrays.

By showing that altered linker spacing is sufficient to change the DNA topology in circular minichromosomes, our results provide a decisive proof to the notion that the chromatin higher-order structure is defined by nucleosome rotational settings (40, 41). Previous crystallographic and EM studies of Richmond and colleagues (32, 33) together with the high-resolution Cryo-EM data (34) revealed the two-start zigzag organization of chromatin fibers for the linkers belonging to the {10*n*} series (L = 20, 30 and 40 bp). Overall, the DNA folding in these fibers remains the same, the main difference being gradual monotonic changes in dimensions of fibers due to increase in linker length. By contrast, here we observe an abrupt change in DNA topology when comparing the fibers with L=20 and 25 bp, consistent with the folding trajectories of DNA linkers being strikingly different in the two cases (Fig. 1). The topological polymorphism of a chromatin fiber depending on linker DNA trajectory, first noted by Crick (15), is reflected in the early studies by Worcel et al. (28), Woodcock et al. (29), and Williams et al. (30) who presented space-filling models of the chromatin fiber with the DNA linking number Δ*Lk* varying from −1 to −2 per nucleosome, depending on the DNA trajectory.

The increased fiber ‘plasticity’ observed for the {10*n*+5} linkers (40-42) is biologically relevant because the {10*n*+5} values are frequently found *in vivo* (36-40). On the other hand, the more tightly folded {10*n*} structures appear to facilitate nucleosome disk stacking by interactions between histone H4 N-terminal domain and histone H2A/H2B acidic patch at the nucleosomal interface (32) and make the structure to be especially sensitive to effects of histone H4 acetylation. Previously it had been shown that histone H4 acetylation imposes a strong unfolding effect on a nucleosome array with L=30 bp (49). In this respect, it would be important to conduct comparative structural and topological studies of {10*n*} and {10*n*+5} chromatin reconstituted with acetylated and nonacetylated histone H4. Our modeling (Fig. 1) predicts that the effect of histone H4K16 acetylation on chromatin structure and DNA topology would be more pronounced for {10*n*} nucleosome spacing.

The experimentally observed topological polymorphism and an extrapolation of our model to include a wide range of variations of linker lengths provides a simple explanation for the observed DNA linking number Δ*Lk* ~ −1.0 in native SV40 minichromosomes (17, 18). These minichromosomes do not have a strong nucleosome positioning and thus may give rise to Δ*Lk* values widely distributed between −0.7 and −1.5 per nucleosome. For nucleosome linkers length close to 60 bp, the most common value for eukaryotic chromatin, the mean value of the Δ*Lk* is about −1.0 per nucleosome (Supplementary Fig. S6). Earlier analysis of distribution of Δ*Lk* for a nucleosome array with randomized nucleosome orientation (27) also showed the Δ*Lk* values close to the −1 per nucleosome.

In living cells, the DNA-dependent processes such as transcription constantly generate positive and negative local DNA supercoiling in accord with the twin-domain model of transcription (50). Because topoisomerases I and II do not completely remove this dynamically induced supercoiling, transcriptionally-active chromosomal domains accumulate regions with the higher and lower superhelical density, which can be detected by a combination of psoralen crosslinking with genome-wide sequencing (9, 51, 52). Recent genome-wide mapping with psoralen revealed extended (on average 100 kb) overtwisted domains correlated with transcriptional activity and chromatin decondensation (10). On the contrary, transcriptionally-silent domains in yeast acquire negative supercoiling (53).

As follows from our findings, the level of DNA supercoiling in chromatin is closely tied with the nucleosome spacing, the supercoiling being more negative (Δ*Lk* ≈ −1.5) and the overall structure more compact (Fig. 1) for the linkers belonging to the {10*n*} series. Apparently, in the yeast genome, the distribution of the nucleosome spacing is uneven and the lowly transcribed genes are characterized by L=10*n*, while the actively transcribed genes predominately have linkers L=10*n*+5 (42). We believe that our findings reflect a universal nature of chromatin structure adapted to the needs of rendering the active chromatin open for transcription by acquiring special nucleosome spacing. More specifically, we propose that by altering nucleosome spacing, transcription may impose slowly-relaxing topological changes on the nucleosome arrays and thus generate topologically and structurally distinct chromosomal domains that will maintain their active and repressed states. In the future, high resolution mapping of nucleosome positioning, which has recently been used to reveal the predominant L=10*n*+5 spacing in the open chromatin of embryonic stem cells (39) may be coupled with structural genome-wide studies probing DNA supercoiling *in situ* (9, 10) to determine the link between nucleosome positioning and DNA topology in complex genomes of higher eukaryotes.

## Methods

### Preparation of circular DNA constructs

The plasmid-based DNA circles have 12 repeats of 601 nucleosome positioning sequence with repeat length of either 167 or 172 bp inserted into pUC19 vector (40). For the pUC19-167x12 plasmid, extra 60 bp were added to make it the same DNA length as the pUC19-172x12 plasmid. Two self-complementary ssDNA fragments, purchased from IDT, were annealed, so that the resulting dsDNA had single stranded overhangs with XbaI site on one end and EcoRI site on the other. The pUC19-167x12 construct was digested with EcoRI and XbaI restriction enzymes and ligated with dsDNA with overhangs using standard procedures. Ligation products were used to transform Stbl2 competent cells; the transformants were grown and plasmid DNA was prepared using Plasmid purification kit (Qiagen) according to the manufacturer’s instructions.

DNA mini-circles containing only 12-mer of 601 nucleosomes with repeat length of 167 and 172 bp were prepared by ligations of linear DNA fragments at low concentration to ensure circle formation rather than ligation of multiple linear fragments. The (167x12)+60 bp and 172x12 bp linear DNA fragments were prepared by cutting off DNA fragments between XbaI and SpeI restriction sites from pUC19 plasmids with corresponding inserts. The DNA fragments were run on 1% agarose gel, bands with inserts were isolated, DNA was extracted with Promega Wizard SV Gel and PCR Clean Up System, cleaned with phenol/chloroform and precipitated. Linear DNA fragments were suspended in 10 mM Tris HCl, pH 7.5. For ligation reaction, DNA was diluted to 1 ng/μl in total volume of 800 μl in multiple tubes. 3200 units of T4 ligase (New England Biolabs) were added to each tube. Reaction was kept at 16 °C for about 48 hours. Every 2-3 hours additional 800 ng of DNA were added to each reaction mixture. Ligation mixtures were concentrated and washed in 10 mM Tris HCL pH 7.5 in Amicon Ultra-4 Ultracel Centrifugal Filters units with MWCO 100 K.

To prepare supercoiled circular DNA (scDNA) with defined superhelical densities (46), 3 μg of DNA samples were treated with various concentrations of ethidium bromide (EtBr) in the presence of 10 units of human Topoisomerase I (TopoGen, cat # TG2005H) in TopoI buffer (10 mM Tris-HCl, pH 7.9, 1 mM EDTA, 150 mM NaCl, 0.1% BSA, 0.1 mM spermidine, 5% glycerol). Mixtures were kept at 37 °C for 2 hours. The reaction was stopped by addition of Na-acetate to 0.33 M and samples were cleaned with phenol/chloroform. After precipitation, scDNA was suspended in 10 mM Tris HCl, pH 7.5.

### Preparation of circular nucleosomal arrays

Nucleosomal arrays were reconstituted by mixing purified chicken erythrocyte core histones (54, 55) with circular DNA. The reconstitution was done as described (40) by a salt dialysis from 2.0 NaCl to 500 mM NaCl followed by dialysis to 10 mM NaCl, 10 mM Tris HCl, pH 8.0, 0.25 mM EDTA.

For TopoI treatment, 1 μg of nucleosomal arrays were mixed with 10 units of Topoisomerase I in TopoI buffer in 50 μl reaction volume for 30 min at 37 °C. The core histones were then removed by SDS/proteinase K treatment, DNA was cleaned with Promega Wizard SV Gel and PCR Clean Up System. DNA samples were run on 1% agarose gels containing different concentrations of CQ in TAE buffer.

**2D electrophoresis** gels were run in 1% agarose at 3 V/cm for 18 h in TAE buffer with CQ concentration as indicated. The gels were removed, soaked for 3 h in TAE buffer with appropriate CQ concentration, placed into e/f apparatus perpendicular to the first direction, and run again for 18 h at 3 V/cm. Gels were stained in GelRed stain (Biotium, cat# 41002) and visualized using Bio-Rad Image Lab imager.

### Transmission Electron Microscopy of nucleosomal arrays

For EM, the reconstituted and Topo I-treated nucleosome arrays were dialyzed in 10 mM HEPES, 10 mM NaCl, 0.1 mM EDTA buffer for 4 hours and then fixed with 0.1% gluteraldehyde overnight at +4 °C. Gluteraldehyde was then removed by dialysis in the same buffer (without glutaraldehyde). Samples were diluted with 50 mM NaCl to ~1 μg/ml, applied to carbon-coated and glow discharged EM grids (T1000-Cu, Electron Microscopy Science) and stained with 0.04% aqueous uranyl acetate as previously described (40). Dark-field images were obtained and digitally recorded as described using JEM-1400 electron microscope (JEOL USA, Peabody, MA) at 120 kV with SC1000 ORIUS 11 megapixel CCD camera (Gatan Inc. Warrendale, PA). The number of nucleosomes formed in each array was counted as described (47).

## Supplementary Figures

**Fig. S1.**
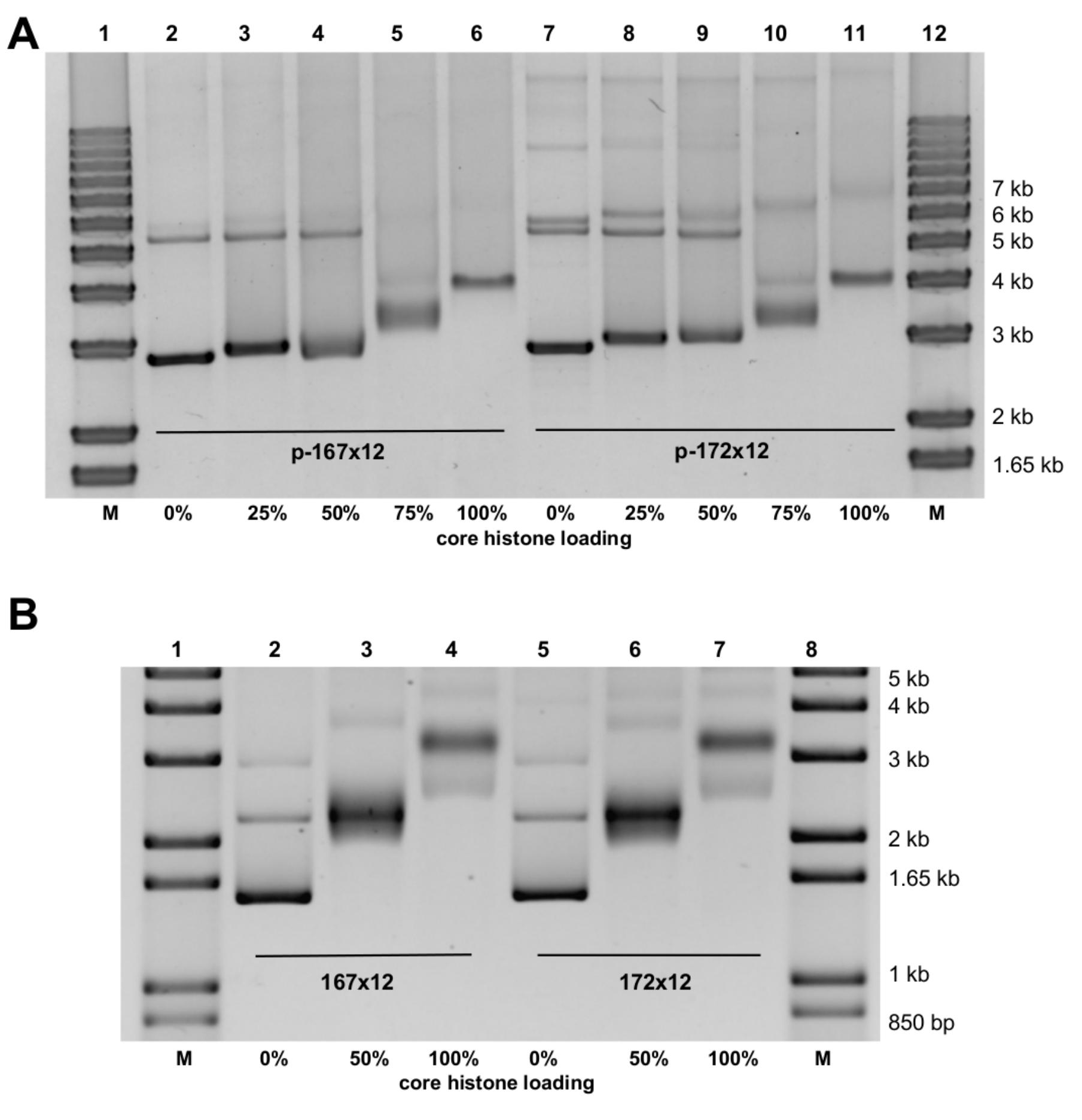
Analysis of circular nucleosome array reconstitution by electrophoretic mobility shift assay. Circular nucleosomal arrays were reconstituted with different core histone loadings and separated on 0.8% type IV agarose DNP gel in TAE buffer. A: plasmid-based circular DNA templates p-167x12 (lanes 2-6) and p-172x12 (lanes 7-11) were reconstituted with 0%, 25%, 50%, 75%, and 100% core histones. B: DNA mini-circular templates 167x12 (lanes 2-4) and 172x12 (lanes 5-7) were reconstituted with 0%, 50% and 100% core histones. M: molecular weight markers.

**Fig. S2.**
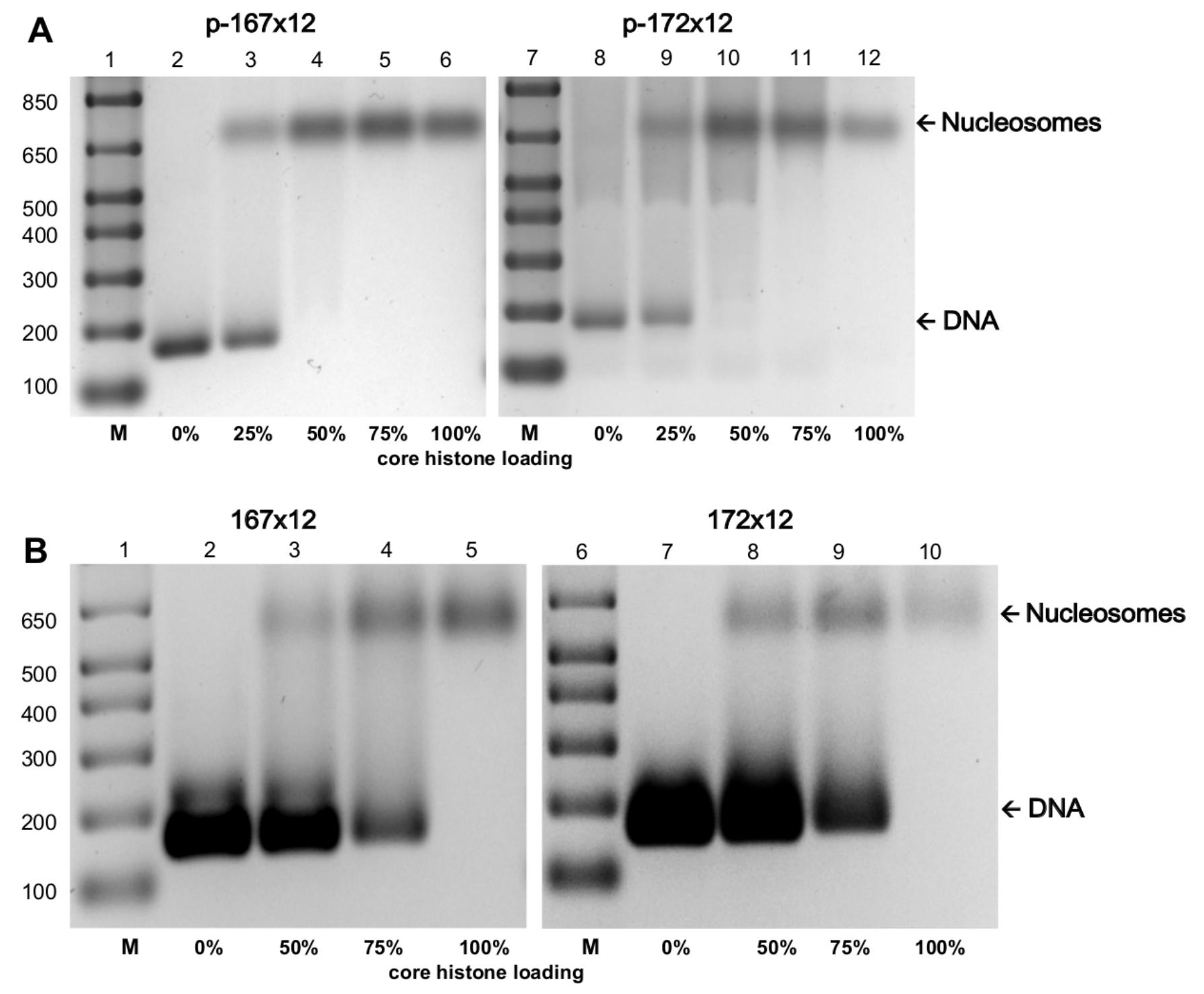
Analysis of circular nucleosome array reconstitution by restriction enzyme digestion and electrophoretic DNA band-shift assay. A: plasmid-based circular DNA templates p-167x12 (lanes 2-6) and p-172x12 (lanes 8-12) were reconstituted with 0%, 25%, 50%, 75%, and 100% core histones and digested with restriction enzymes EcoRV and SpeI (167x12) and XbaI, SpeI, and NlaIII (172x12) to cut off the vector and excise the mononucleosomes. B: DNA mini-circular templates 167x12 (lanes 2-5) and 172x12 (lanes 7-10) were reconstituted with 0%, 50%, 75%, and 100% core histones and digested with restriction enzymes EcoRV (167x12) and NlaIII (172x12) to excise the mononucleosomes. The free DNA and histone-associated mononucleosomes were separated on native 1.5% type IV agarose gel in TAE buffer. M: molecular weight markers.

**Fig. S3.**
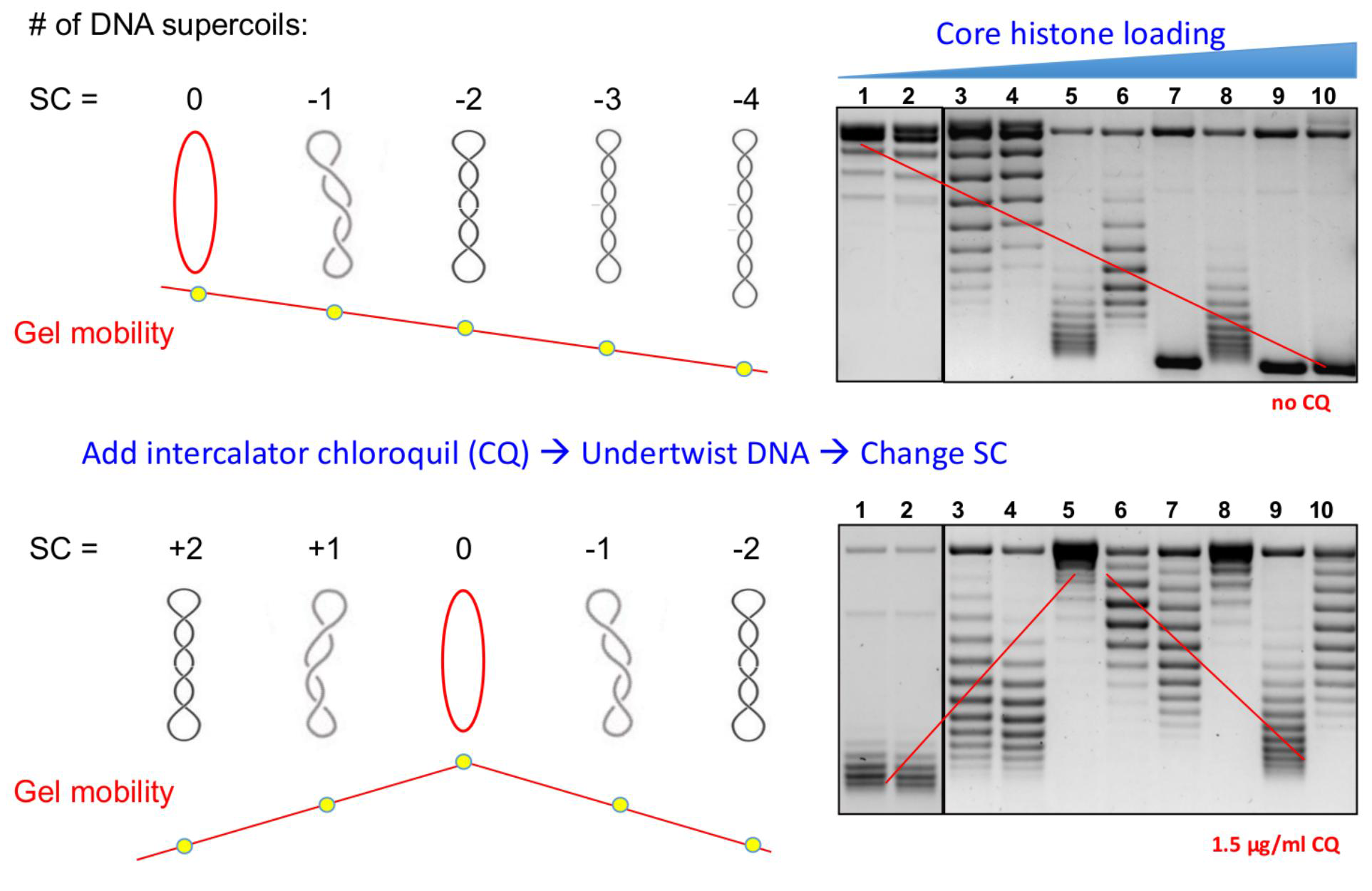
Binding of intercalator CQ to supercoiled DNA changes its electrophoretic mobility. **Top**: In the absence of intercalator, the topoisomers with a higher absolute number of superhelical turns, |SC|, move faster in agarose gel. **Bottom:** Binding of intercalator partially unwinds the DNA duplex (i.e., its Twist is decreased). Accordingly, the Writhe is increased (because the sum of Twist and Writhe remains unchanged in circular DNA), which corresponds to the increase in the number of DNA supercoils, SC. This, in turn, leads to a change in the gel mobility of DNA. For example, the topoisomer shown on the left becomes positively supercoiled (SC=2) and moves down relatively fast (compared to its low mobility in the absence of intercalator). By contrast, binding of intercalator to the topoisomer in the center compensates its negative supercoils and it runs at the top of gel as relaxed DNA with SC=0. For the rightmost topoisomer, the SC value changes from −4 to −2; it still migrates in gel relatively fast (but not as fast as in the absence of intercalator). Note that this simple scheme accounts for the mobility of scDNA in agarose gel – compare the red lines on the left with the lines in the right panels highlighting mobility of the dominant DNA topoisomers in the gels with [CQ]=0 and 1.5 μg/ml (see Fig. 2 A and Fig. 2 B). The core histones loading increases from zero to 100%: lanes 1 and 2, 0%; lanes 3 and 4, 25%; lanes 5 and 6, 50%; lanes 7 and 8, 75%; lanes 9 and 10, 100%. The odd and even lanes show gel mobility of the p-167x12 and p-172x12 constructs, respectively.

**Fig. S4.**
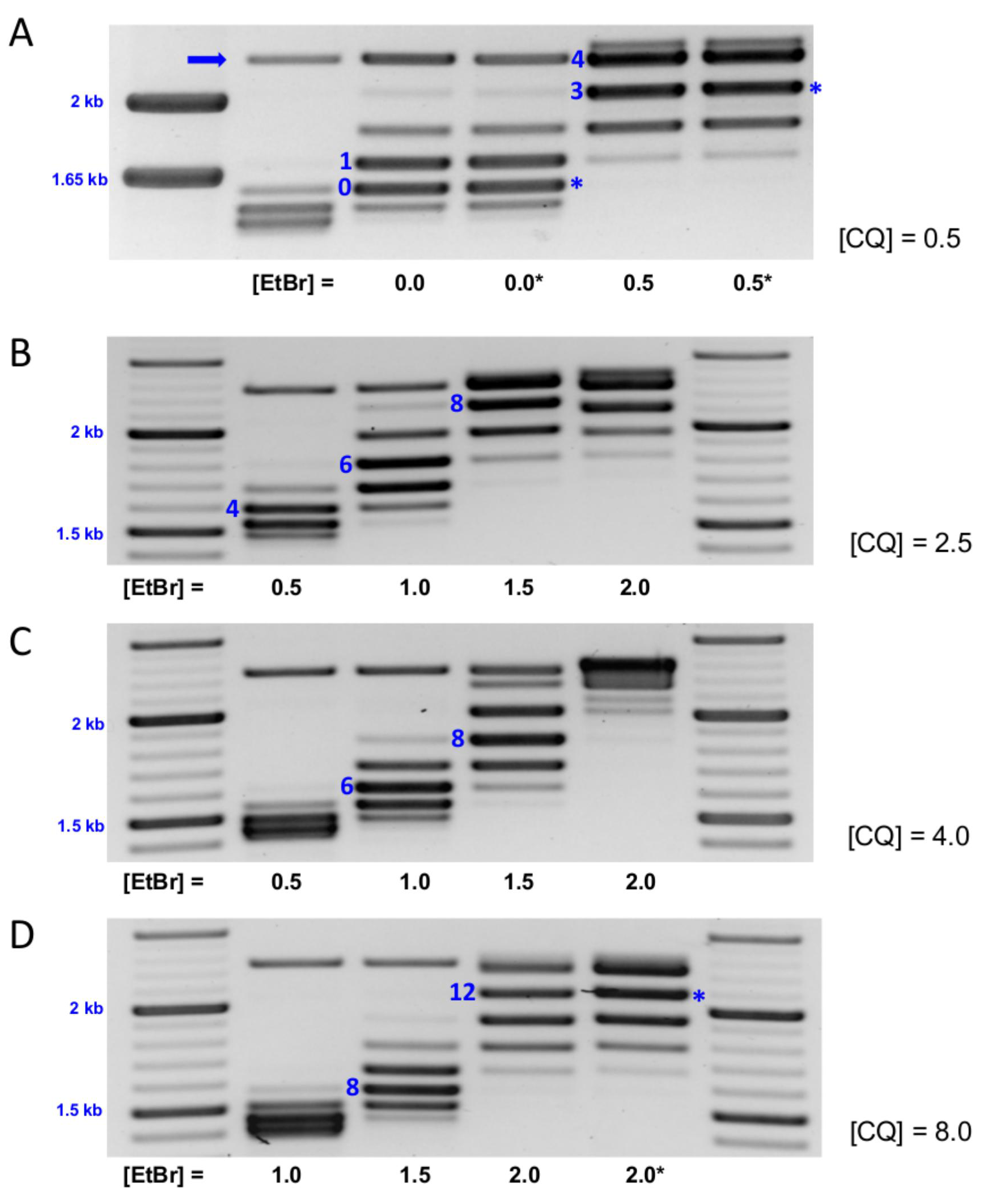
Modulating negative supercoiling in DNA mini-circles in the presence of various concentrations of EtBr. Vector-free circular DNA templates were treated with Topoisomerase I in the presence of 0.5 – 2.0 μg/ml EtBr as indicated at the bottom of the gels and separated on agarose gels in the presence of 0.5 (A), 2.5 (B), 4.0 (C), and 8.0 (D) μg/ml CQ in the gel and TAE buffer. The data are presented for the 167x12 construct, except where the bands (and the [EtBr]) values) are marked with asterisks (A and D). In the latter cases the 172x12 construct was used. The left lane (A) and the left and right lanes (B-D) contain molecular markers. The second lane (A) contains ligated 172x12 construct (without TopoI relaxation). The position of form II (nicked) circles is indicated by arrow. The numbers in blue indicate the absolute value of the DNA linking number, |Δ*Lk*|, measured relative to the ‘relaxed’ topoisomer marked 0 in (A). For the lanes with [EtBr]=0: note that the 167x12 and 172x12 constructs have equal distribution of DNA topoisomers obtained after relaxation by TopoI. Also note that the scDNA obtained at [EtBr]=2 μg/ml in the presence of 8.0 μg/ml CQ has the strongest band corresponding to Δ*Lk* = −12 (D).

**Fig. S5.**
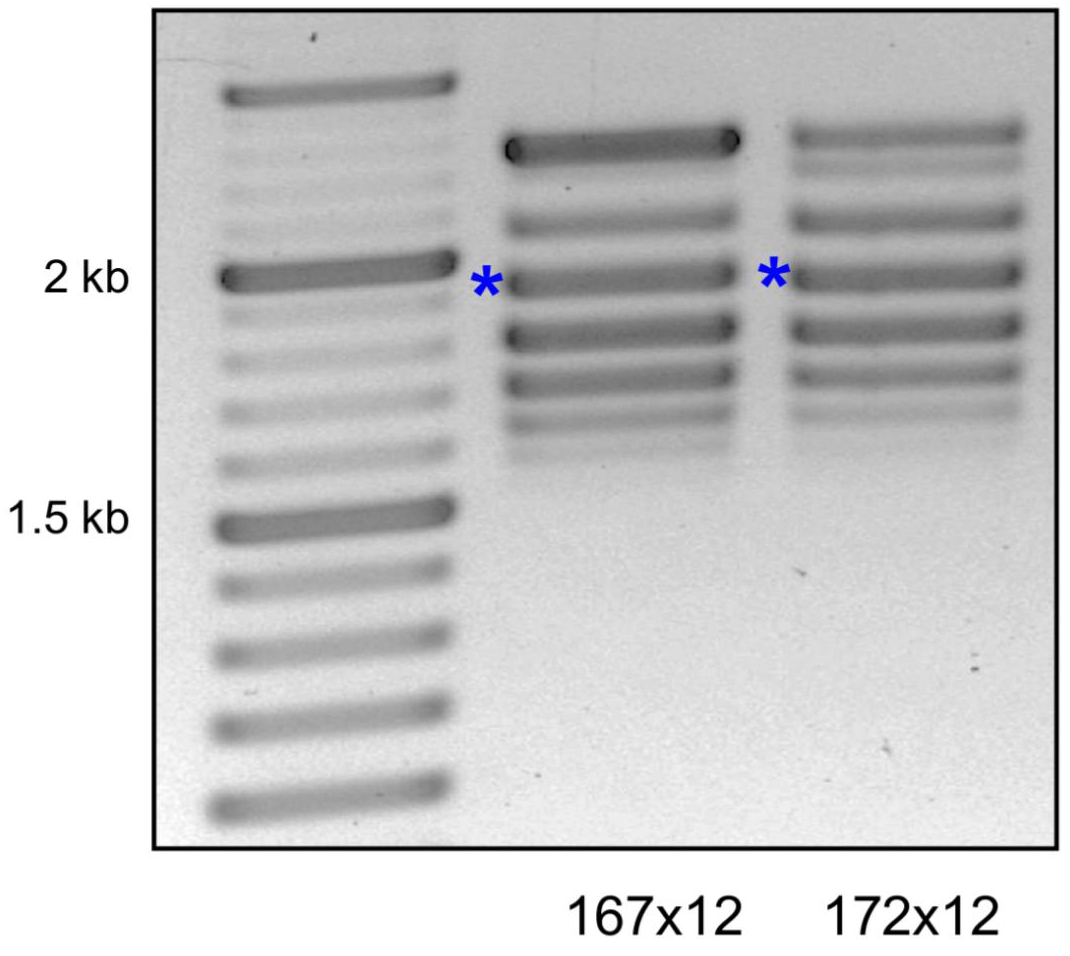
Preparation of supercoiled DNA constructs 167x12 and 172x12 relaxed by TopoI in the presence of 4.0 μg/ml EtBr. Note that the two mini-circles have practically the same distribution of superhelical topoisomers (the strongest bands are marked by asterisks). Agarose gel was run in the presence of 8 μg/ml CQ.

**Fig. S6.**
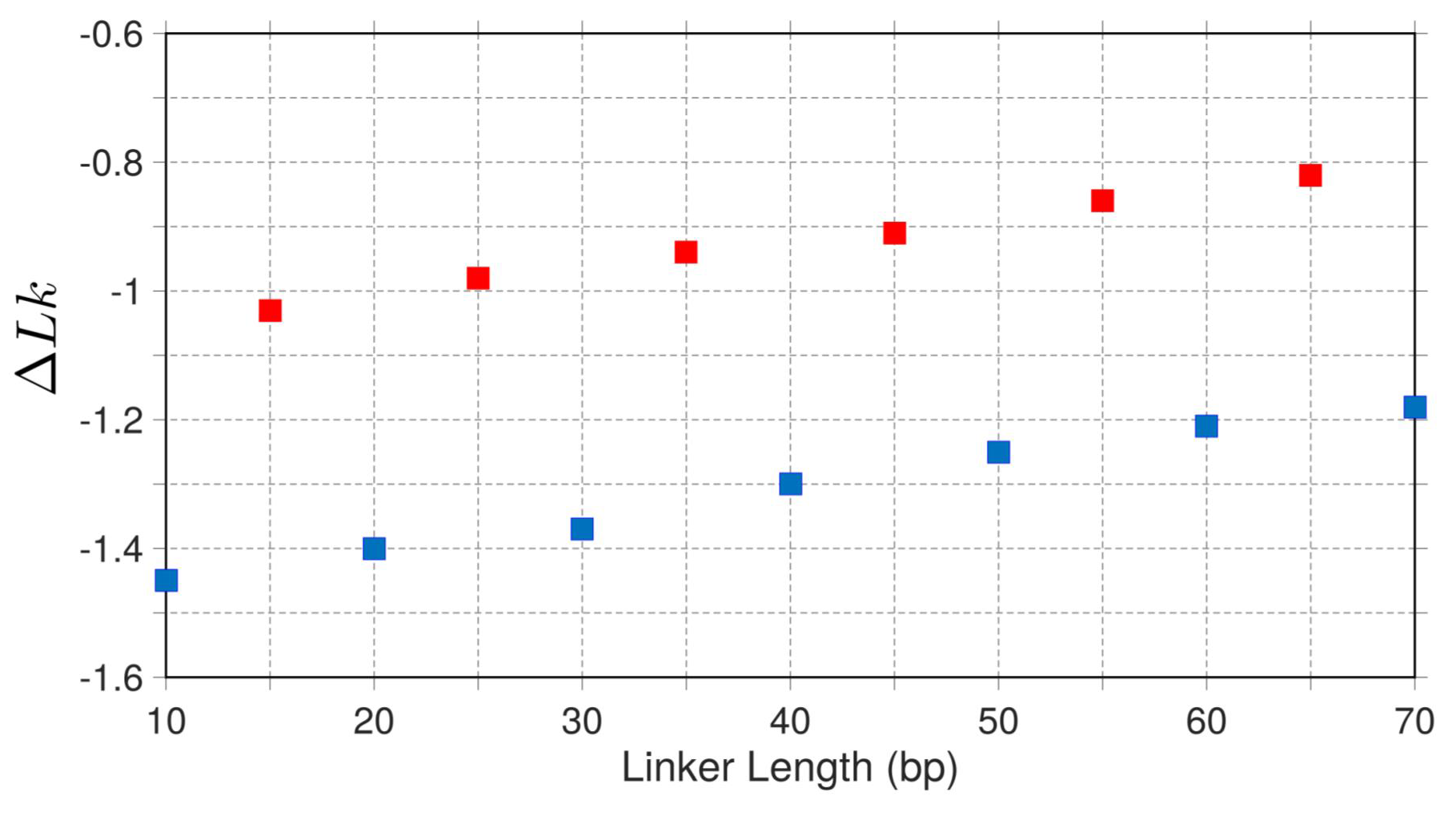
Linking number per nucleosome, Δ*Lk*, in regular fibers with various linker lengths. The Δ*Lk* values calculated for arrays of 100 nucleosomes (42), are shown for the two series, {L = 10*n*} (blue squares) and {L = 10*n* + 5} (red squares).

